# A putative antiviral role of plant cytidine deaminases

**DOI:** 10.1101/005256

**Authors:** Susana Martín, José M. Cuevas, Ana Grande-Pérez, Santiago F. Elena

**Affiliations:** Instituto de Biología Molecular y Celular de Plantas, Consejo Superior de Investigaciones Científicas-Universidad Politécnica de Valencia, Campus UPV CPI 8E, Ingeniero Fausto Elio s/n, 46022 València, Spain; Instituto de Hortofruticultura Subtropical y Mediterránea “La Mayora”, Consejo Superior de Investigaciones Científicas-Universidad de Málaga, Campus de Teatinos, 29071 Málaga, Spain; Área de Genética, Universidad de Málaga, Campus de Teatinos, 29071 Málaga, Spain; The Santa Fe Institute, 1399 Hyde Park Road, Santa Fe NM 87501, USA

**Author notes:** Correspondence and requests for materials should be addressed to S.F.E. These authors contributed equally to this work (Susana Martín, José M. Cuevas).

## Abstract

A mechanism of innate antiviral immunity operating against viruses infecting mammalian cells has been described during the last decade. Host cytidine deaminases (e.g., APOBEC3 proteins) edit viral genomes giving raise to hypermutated nonfunctional viruses; consequently, viral fitness is reduced through lethal mutagenesis. By contrast, sub-lethal hypermutagenesis may contribute to virus evolvability by increasing population diversity. To prevent genome editing, some viruses have evolved proteins that mediate APOBEC3 degradation. The model plant *Arabidopsis thaliana* encodes for nine cytidine deaminases (AtCDAs), raising the question of whether deamination is an antiviral mechanism in plants as well. Here we tested the effects of AtCDAs expression on the pararetrovirus *Cauliflower mosaic virus* (CaMV). We show that *A. thaliana AtCDA1* gene product exerts a mutagenic activity, which indeed generates a negative correlation between the level of *AtCDA1* expression and CaMV accumulation in the plant, suggesting that deamination may also work as an antiviral mechanism in plants.

---

Human APOBEC family includes enzymes that catalyze the hydrolytic deamination of cytidine to uridine or deoxycytidine to deoxyuridine. This family is composed by eleven known members: APOBEC1, APOBEC2, APOBEC3 (further classified as A3A to A3H), APOBEC4, and AID (activation induced deaminase). APOBEC proteins are associated to several functions involving editing of DNA or RNA (reviewed in ref. 1). APOBEC1 (apolipoprotein B editing catalytic subunit 1) mediates deamination of cytidine at position 6666 of apolipoprotein B mRNA, resulting in the introduction of premature stop codon and the production of the short form of the protein^2–4^. APOBEC2 is essential for muscle tissue development^5^. APOBEC4 has no ascribed function so far^6^. AID deaminates genomic ssDNA of B cells, initiating immunoglobulin somatic hypermutation and class switch processes^7–9^. Most notably, APOBEC3 enzymes participate in innate immunity against retroviruses and endogenous retroelements^10–12^. Sheehy *et al.* demonstrated that A3G also plays a role in immunity against *Human immunodeficiency virus* type 1 (HIV-1)^13^. For its antiviral role A3G is packaged along with viral RNA^14^. Upon infection of target cells and during the reverse transcription process, A3G deaminates the cytosine residues of the nascent first retroviral DNA strand into uraciles. The resulting uracil residues serve as templates for the incorporation of adenine, which at the end result in strand-specific C/G to T/A transitions and loss of infectivity through lethal mutagenesis^15–19^. At the other hand, a sub-lethal mutagenic activity of APOBEC3 proteins may contribute as an additional source for HIV-1 genetic diversity, hence bolstering its evolvability^20–22^. APOBEC3 proteins had been shown to inhibit other retroviruses (*Simian immunodeficiency virus*^23^, *Equine infectious anemia virus*^24^, *Foamy virus*^25^, *Human T-cell leukemia virus*^26^, and *Murine leukemia virus*^27^), pararetroviruses (*Hepatitis B virus*^28^) and DNA viruses (*Herpes simplex virus* 1 (HSV-1)^29, 30^ and *Epstein-Barr virus* (EBV)^30^). In the cases of HSV-1 and EBV antiviral role of deaminase has not been demonstrated^30^. Evidences also exist that A3G significantly interferes with negative-strand RNA viruses lacking a DNA replicative phase^31^. For example, the transcription and protein accumulation of *Measles virus*, *Mumps virus* and *Respiratory syncytial virus* was reduced 50 to 70% whereas the frequency of C/G to U/A mutations was ∼4-fold increased^31^. In contrast, A3G plays no antiviral activity against *Influenza A virus* despite being highly induced in infected cells as part of a general IFNβ response to infection^32, 33^.

Human APOBEC belongs to a superfamily of polynucleotide cytidine and deoxycytidine deaminases distributed throughout the biological world^34^. All family members contain a zinc finger domain (CDD), identifiable by the signature (H/C)-x-E-x25-30P-C-x-x-C. Plants are not an exception and, for example, *Arabidopsis thaliana* encodes for nine putative cytidine deaminases (named as *AtCDA1* to *AtCDA9* genes). Whilst the *AtCDA1* gene is located in chromosome II, the other eight genes are located in chromosome IV. In the case of rice and other monocots, only a CDA has been identified^34^. Interestingly, this CDA expression was highly induced as part of the general stress response of rice against the infection of the fungal pathogen *Magnaporthe grisea*, resulting in an excess of A to G and U to C mutations in defense-related genes^35^. Edited dsRNAs might be retained in the nucleus and degraded, generating miRNAs and siRNAs^36^. Provided the presence of the catalytic deamination motif in AtCDA proteins and the relevance of deamination as an antiviral innate response in animals, we sought first to determine whether any of the AtCDAs can participate in deaminating the genome of the pararetrovirus *Cauliflower mosaic virus* (CaMV; *Caulimovirus*, *Caulimoviridae*) and, second, to explore whether this deamination may negatively impact viral infection. We hypothesize that deamination may take place mainly at the reverse transcription step. CaMV genome is constituted by a single molecule of circular double-stranded DNA of 8 kpb^37^. The DNA of CaMV has three discontinuities, Δ1 in the minus-sense strand (or *a* strand) and Δ2 and Δ3 in the plus-sense strand (yielding the *b* and *g* strands). In short, the replication cycle of CaMV is as follows^37^: in the nucleus of the infected cell, the *a* strand is transcribed into a 35S RNA, with terminal repeats, that migrates to the cytoplasm. Priming of the 35S RNA occurs by the annealing of the 3' end of tRNA^met^ to the primer-binding site (PBS) sequence, leading to the synthesis of the DNA *a* strand by the virus' reverse transcriptase. Then, the RNA in the heteroduplex is degraded by the virus' RNaseH activity, leaving purine-rich regions that act as primers for the synthesis of the plus-sense DNA *b* and *g* strands.

Our results show that *AtCDA1* significantly increases the number of G to A mutations *in vivo*, and that there is a negative correlation between the amount of *AtCDA1* mRNA present in the cell and the load reached by CaMV, suggesting that deamination of viral genomes may also constitute a significant antiviral mechanism in plants.

## Results

### Effect of AtCDAs overexpression on CaMV mutational spectrum

To test the mutagenic activity of *A. thaliana* CDAs, nine *Nicotiana bigelovii* plants were inoculated with CaMV. After systemic infection was established, we performed transitory AtCDA overexpression experiments. To do so, the same leaf was agroinfiltrated twice; half leaf was infiltrated with one of the nine AtCDA genes and the other half leaf was infiltrated with the empty vector. This test was done for all nine AtCDA genes in different plants. The presence of AtCDA mRNAs was verified by RT-PCR from DNase-treated RNA extracts. DNA was extracted from agroinfiltrated areas for 3D-PCR amplification of a 229 bp fragment in the ORF VII of CaMV. 3D-PCR uses a gradient of low denaturation temperatures during PCR to identify the lowest one, which potentially allows differential amplification of A/T rich hypermutated genomes^38^. There were no differences in the lowest denaturation temperature resulting in amplification between the controls and the AtCDA-agroinfiltrated samples, suggesting that hypermutated genomes should be at low frequency, if present at all, in the latter case.

PCR products obtained at the lowest denaturation temperature were cloned and sequenced. In a preliminary experiment we sequenced 25 clones from each AtCDA/negative control pair (Supplementary Table S1). At least one G to A transition was detected in clones from areas infiltrated with *AtCDA1*, *AtCDA2* and *AtCDA9* genes. For these three genes, we further increased the number of sequenced clones up to 106. The CaMV mutant spectra was significantly different between plants overexpressing *AtCDA1* and their respective negative controls (Fig. 1a: χ^2^ = 25.760, 7 d.f., *P* = 0.001), being the difference entirely driven by the 471.43% increase in G to A transitions observed in the plants overexpressing *AtCDA1*. A thorough inspection of alignments showed that most of the G to A mutations (65.6%) detected in the different samples were located at the nucleotide position 181 (Supplementary Table S1). By contrast, no overall difference existed between the mutant spectra of CaMV populations replicating in plants overexpressing *AtCDA2* (Fig. 1b: χ^2^ = 8.944, 6 d.f., *P* = 0.177) or *AtCDA9* genes (Fig. 1c: χ^2^ = 6.539, 8 d.f., *P* = 0.587) and their respective controls. Consistently, the mutant spectra from the three AtCDA-overexpressed samples were significantly heterogeneous (χ^2^ = 41.063, 16 d.f., *P* = 0.001), again due to the enrichment in G to A transitions observed in the case of *AtCDA1*. By contrast, the three independent control inoculation experiments showed homogeneous mutant spectra for CaMV (χ^2^ = 14.605, 18 d.f., *P* = 0.689) and undistinguishable from the mutant spectra previously reported for natural isolates of this virus^39^. This later comparison confirms the stability of CaMV mutant spectrum when not perturbed by e.g. host mutagenic enzymes and the host species is always the same.

**Figure 1.**
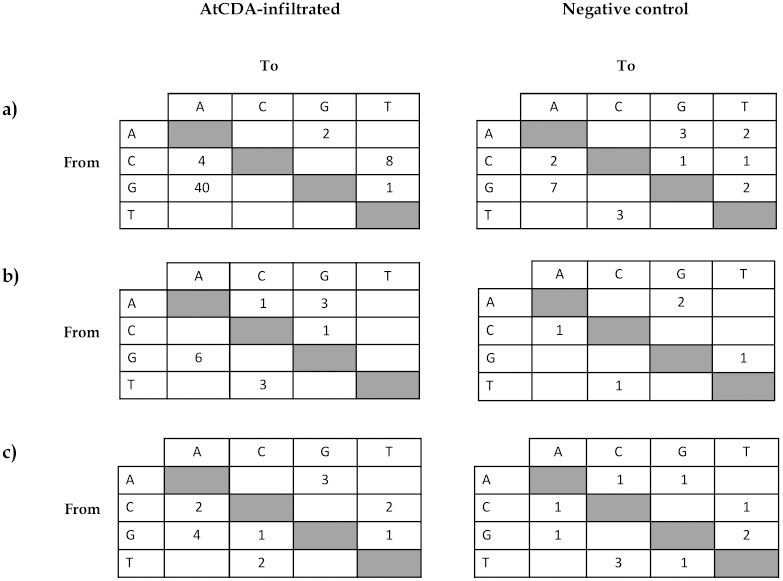
Number of mutations in CaMV genomes isolated from plant tissues agroinfiltrated with different CDAs. (A) *AtCDA1*, (B) *AtCDA2* and (C) *AtCDA9*. The pBIN61 empty vector was agroinfiltrated in the same leaves than their corresponding CDAs (mock). For each sample 20034 nucleotides were sequenced.

Therefore, we conclude that overexpressing the *AtCDA1* gene resulted in a significant shift in CaMV genome composition towards G to A mutations, as expected from a cytidine deaminase hypermutagenic activity.

### Effect of suppressing AtCDA expression on the viral load and mutational spectrum of CaMV

To test the effects of suppressing the expression of AtCDA on viral accumulation we produced a transgenic line of *A. thaliana* Col-0, named miR1-6-3. This line was stably transformed with an artificial micro-RNA (amiR), controlled by the *Aspergillus nidulans* ethanol regulon to achieve ethanol-triggered RNAi-mediated simultaneous suppression of AtCDAs 1, 2, 3, 4, 7, and 8 expression. Transgenic and wild-type plants were subjected to periodical treatment with 2% ethanol (or water for the control groups). Subsequently, plants were inoculated with the infectious clone pCaMVW260 that expresses the genome of CaMV. Samples were taken eight days after inoculation and *AtCDA1* mRNA and CaMV viral load were quantified by real time RT-qPCR and qPCR, respectively, in the same samples. For each genotype and/or treatment, 22 plants were analyzed.

The expression of *AtCDA1* mRNA depended on the plant genotype (Fig. 2a; GLM: χ^2^ = 28.085, 1 d.f., *P* < 0.001) as well as on the interaction of plant genotype and treatment (χ^2^ = 26.037, 1 d.f., *P* < 0.001), suggesting a differential accumulation of *AtCDA1* mRNA on each plant genotype depending on the amiR induction state. Ethanol treatment reduced the amount of *AtCDA1* mRNA by 24.01% in transgenic plants, proving that triggering the expression of the amiR significantly and efficiently silences the expression of *AtCDA1*. Unexpectedly, the effect was the opposite in wild-type plants, for which we observed 23.76% increase in *AtCDA1* mRNA accumulation (Fig. 2a) upon treatment with ethanol. This increase in the expression of *AtCDA1* in wild-type plants after the ethanol treatment is certainly puzzling and the underlying mechanisms deserve investigation by its own. However, for the purpose of this study, its relevance is that it may increase the number of G to A mutations in CaMV genome, thus making the antiviral effect stronger to some extent.

**Figure 2.**
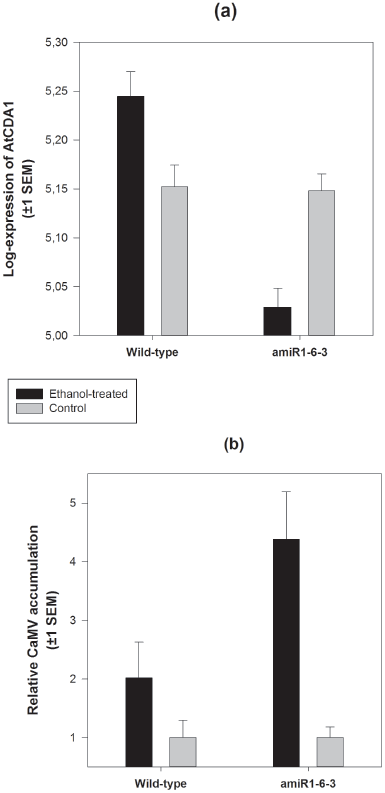
Accumulation of *CDA1* mRNA molecules and CaMV genomes. **(a)** Number of *AtCDA1* mRNA molecules/80 ng total RNA quantified by RT-qPCR using the standard curve method for absolute quantification. **(b)** Number of CaMV genomes/80 ng total DNA. For each block of plants (wild-type and amiR1-6-3), values were normalized to the average number of genomes estimated in the corresponding water-treated (control) plants.

More interestingly, the relative accumulation of CaMV in ethanol-treated plants was significantly different depending on the plant genotype being infected (Fig. 2b; Mann-Whitney *U* test, *P* = 0.002): silencing the *AtCDA1* gene bolstered CaMV accumulation in 103.10% compared to the accumulation observed in wild-type plants. Furthermore, a significant negative correlation between the number of molecules of *AtCDA1* mRNA and viral load was detected (partial correlation coefficient controlling for treatment: *r* = −0.299, 86 d.f., *P* = 0.005).

Provided the significant increase of viral load in plants with lower levels of *AtCDA1* mRNA, we sought for the molecular signature of deamination in transgenic plants. For this, we selected three biological replicates from each treatment group (ethanol or control) and sequenced (as previously explained) from 39 to 45 clones of the CaMV fragment from each replicate. As shown in Fig. 3, silencing of the *AtCDA1* gene affects the composition of CaMV mutant spectrum by reducing the number of G to A transitions in 69.23%. Nonetheless, despite this reduction goes in the expected direction, overall, both mutational spectra were not significantly different (Fig. 3: χ^2^ = 9.108, 6 d.f., *P* = 0.168), prompting caution against making a strong conclusion on the role of deamination in the observed increase in CaMV accumulation.

**Figure 3.**
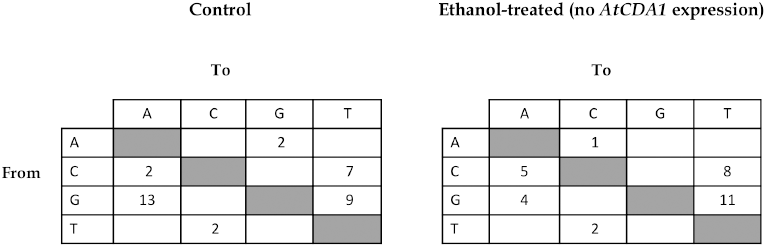
Number of mutations found pooling the CaMV sequences from ethanol-treated and control amiR1-6-3 plants (3 replicates). The number of nucleotides sequenced was 23436 for control and 24003 for ethanol-treated plants. Ethanol-treated plants turn on the expression of amiR1-6-3 that was designed to silence the expression of the *AtCDA1* gene.

Therefore, we conclude that suppressing simultaneously the expression of the AtCDAs 1, 2, 3, 4, 7, and 8 significantly reduces the accumulation of CaMV. However, the characterization of the mutant spectrum of the same CaMV populations provides no strong enough support to the cytidine deamination hypothesis.

## Discussion

Lethal mutagenesis through deamination of RNA/DNA by cytidine deaminases has been proven to work as an antiviral mechanism against retroviruses^16–19, 23–27^, and some DNA^28–30^ and RNA^31^ viruses infecting mammals. Our results show that *A. thaliana CDA1* gene has some degree of mutagenic activity on the pararetrovirus CaMV genome. Moreover, simultaneous suppression of the expression of a subset of AtCDAs, including *AtCDA1*, increased CaMV load, strongly suggesting an antiviral role for AtCDAs.

Our data show that AtCDAs probably restrict CaMV replication through a process similar to the restriction of HIV-1 by APOBEC3. CaMV replicates in the cytoplasm by reverse transcription using the plus-strand 35S RNA as template. As for HIV-1, first strand negative cDNA could be deaminated during reverse transcription, transforming deoxycytidine into deoxyuridine. Then, when the plus strand is produced an adenine is incorporated instead of a guanine, increasing the proportion of G to A mutations. In the case of HIV-1, this G to A mutational bias is explained by A3G and A3H specificity for single negative stranded DNA because during HIV-1 replication. Cytosine to guanine transitions are rare and restricted to the PBS site and U3 region in the 5' long terminal repeat where plus stranded DNA is predicted to become transiently single stranded^40^. Similarly, during CaMV replication the negative strand remains single stranded, while the positive is copied from it and remains double stranded^41^. Surprisingly, for *AtCDA1* C to T mutations were also increased; the region studied here is close to the 5' end of CaMV, which contains the PBS for minus strand synthesis and the ssDNA discontinuity Δ1. The observed C to T transitions could reflect transient positive stranded ssDNA in the 5' terminal region during reverse transcription, nevertheless a different substrate specificity of *A. thaliana* CDAs cannot be ruled out.

Most of the G to A transitions detected in agroinfiltration experiments were located in the G at position 181. HIV-1 hypermutated genomes show mutational hot spots as well, which are due to preference of A3G and A3F for deamination of the third cytidine in 5'-CCC (minus strand) and 5'-GGC, respectively^42, 43^. The context of the cytidine complementary to G181 (5'-GGC) differs from those described for APOBEC3, suggesting that if *A. thaliana* CDAs had context preference, it would be different to A3G. However, the low number of mutations found does not allow concluding a possible context preference for AtCDAs. Since our experiments are performed *in vivo*, negative selection is expected to purge genomes carrying deleterious mutations. This limitation could account for our failure to detect largely hypermutated genomes and makes necessary to develop new selection-free assays to further characterize AtCDA-induced mutagenesis.

Although there is not a demonstrated correlation between the expression of APOBEC3 and mutational bias of viruses infecting mammals, caulimoviruses have an excess of G to A transition in synonymous positions^44^. In *A. thaliana* plants, we found that silencing of *AtCDA1* reduced the frequency of G to A transitions in CaMV genome, suggesting a contribution of AtCDAs to the nucleotide bias found in caulimoviruses. The increased viral load in CDA-silenced *A. thaliana* plants strongly suggest that deamination of viral genomes may work as an antiviral mechanism in plants, opening questions about how general this mechanism might be, and also about what its contribution to viral evolution shall be. Describing a new natural antiviral mechanism in plants opens new research avenues for the development of new durable control strategies.

## Methods

### Transient overexpression of AtCDAs in *N. bigelovii* plants infected with CaMV

AtCDAs cDNAs were cloned under the 35S promoter in pBIN61 vector^45^. *N. bigelovii* plants were inoculated with CaMV virions purified from *Brassica rapa* plants^46^ previously infected with the clone pCaMVW260^47^. Symptomatic leafs were agroinfiltrated^45^ with one of the nine AtCDAs and with the empty vector pBin61, each on one half leaf. Samples were collected three days post-agroinfiltration.

### Inducible cosuppression of multiple AtCDAs expression by RNAi

The design and cloning of the amiR able to simultaneously suppress the expression of AtCDAs 1, 2, 3, 4, 7, and 8 was performed as described in ref. 48. The amiRNA was cloned under the control of *A. nidulans* ethanol regulon^49, 50^ and used to transform *A. thaliana* by the floral dip method^51^. By doing so, we obtained the transgenic line miR1-6-3. One-month-old seedlings of transgenic and wild-type *A. thaliana* were treated with 2% ethanol (or water for the control groups) three times every four days. Three days after the third treatment, plants were inoculated with the infectious clone pCaMVW260 as described in ref. 47. Infections were established by applying 1.31×10^11^ molecules of pCaMVW260 to each of three leaves per plant. Subsequently, plants were subjected to two additional treatments with 2% ethanol (or water) one and five days post-infection. Finally, samples were taken eight days after inoculation and handled as previously described^51^. For each genotype (transgenic or wild-type) and treatment (ethanol or water) combination, 22 plants were analyzed.

### Detection of A/T enriched genomes

CaMV genomic DNA was purified using DNeasy Plant Mini Kit (Qiagen) according to manufacturer's instructions. For detection of edited genomes 3D-PCR was performed using primers HCa8Fdeg and HCa8Rdeg (Supplementary Table 2). PCRs were performed in a Mastercycler® (Eppendorf) at denaturation temperatures 82.1°C, 82.9°C, 83.9°C, and 85.0°C. PCR products obtained with the lowest denaturation temperature were cloned in pUC19 vector (Fermentas), transformed in *Escherichia coli* DH5α and sent to GenoScreen (Lille, France) for sequencing.

### RT-qPCR analysis of *AtCDA1* mRNA and qPCR analysis of CaMV load in transgenic plants

Total RNA was extracted from *A. thaliana* plants using the RNeasy® Plant Mini Kit (Qiagen), according to manufacturer's instructions. *AtCDA1* specific primers qCDA1-F and qCDA1-R were designed using Primer Express software (Applied Biosystems). RT-qPCR reactions were performed using the One Step SYBR PrimeScript RT-PCR Kit II (Takara). Amplification, data acquisition and analysis were carried out using an Applied Biosystems Prism 7500 sequence detection system. All quantifications were performed using the standard curve method. To quantify *AtCDA1* mRNA, a full-ORF runoff transcript was synthetized with T7 RNA polymerase (Roche) using as template a PCR product obtained from cloned AtCDA1 and primers T7-CDA1F and qCDA1-R. CaMV qPCR quantitation was performed as described in ref. 51.

### Primers

All primers used are listed in Supplementary Table 2.

## Acknowledgements

We thank Francisca de la Iglesia, Àngels Pròsper and Ana Cuadrado for excellent technical assistance, Miguel A. Blázquez for help in designing the amiR1-6-3 and generating the transgenic plants and Rémy Froissart for providing the pCaMVW260 infectious clone. This work was supported by the Spanish Ministerio de Ciencia e Innovación grant BFU2009-06993 to S.F.E. J.M.C. was supported by the CSIC JAE-doc program/Fondo Social Europeo. A.G.-P. was supported by a grant for Scientific and Technical Activities and by grant P10-CVI-65651, both from Junta de Andalucía.

## Author contributions

S.F.E. conceived the study, designed the experiments and analyzed the data. S.M., J.M.C. and A.G.-P. performed the experiments and contributed to experimental design. S.M., J.M.C. and S.F.E. wrote the paper. All authors revised and approved the manuscript.

## References

1. Smith, H.C., Bennett, R.P., Kizilyer, A., McDougall, W.M. & Prohaska, K.M. Functions and regulation of the APOBEC family of proteins. Semin. Cell Dev. Biol. 23, 258–268 (2011).

2. Driscoll, D.M. & Zhang, Q. Expression and characterization of p27, the catalytic subunit of the apolipoprotein B mRNA editing enzyme. J. Biol. Chem. 269, 19843–19847 (1994).

3. Navaratnam, N. et al. The p27 catalytic subunit of the apolipoprotein B mRNA editing enzyme is a cytidine deaminase. J. Biol. Chem. 268, 20709– 20712 (1993).

4. Teng, B., Burant, C.F. & Davidson, N.O. Molecular cloning of an apolipoprotein B mRNA editing protein. Science 260, 1816–1819 (1993).

5. Sato, Y. et al. Deficiency in APOBEC2 leads to a shift in muscle fiber type, diminished body mass, and myopathy. J. Biol. Chem. 285, 7111–7118 (2010).

6. Rogozin, I.B., Basu, M.K., Jordan, I.K., Pavlov, Y.I. & Koonin, E.V. APOBEC4, a new member of the AID/APOBEC family of polynucleotide (deoxy)cytidine deaminases predicted by computational analysis. Cell Cycle 4, 1281–1285 (2005).

7. Muramatsu, M. et al. Specific expression of activation-induced cytidine deaminase (AID), a novel member of the RNA-editing deaminase family in germinal center B cells. J. Biol. Chem. 274, 18470–18476 (1999).

8. Arakawa, H., Hauschild, J. & Buerstedde, J.M. Requirement of the activation-induced deaminase (AID) gene for immunoglobulin gene conversion. Science 295, 1301–1306 (2001).

9. Fugmann, S.D. & Schatz, D.G. One AID to unite them all. Science 295, 1244–1245 (2002).

10. Chiu, Y.L. et al. High-molecular-mass APOBEC3G complexes restrict *Alu* retrotransposition. Proc. Natl. Acad. Sci. USA 103, 15588–15593 (2006).

11. Schumann, G.G. APOBEC3 proteins: major players in intracellular defence against LINE1-mediated retrotransposition. Biochem. Soc. Trans. 35, 637–642 (2007).

12. Esnault, C., Millet, J., Schwartz, O. & Heidmann, T. Dual inhibitory effects of APOBEC family proteins on retrotransposition of mammalian endogenous retroviruses. Nucl. Acids Res. 34, 1522–1531 (2006).

13. Sheehy, A.M., Gaddis, N.C., Choi, J.D. & Malim, M.H. Isolation of a human gene that inhibits HIV-1 infection and is suppressed by the viral Vif protein. Nature 418, 646–650 (2002).

14. Smith, H.C. APOBEC3G: a double agent in defense. Trends Biochem. Sci. 36, 239–244 (2011).

15. Mangeat, B. et al. Broad antiretroviral defence by human APOBEC3G through lethal editing of nascent reverse transcripts. Nature 424, 99–103 (2003).

16. Zhang, H. et al. The cytidine deaminase CEM15 induces hypermutation in newly synthesized HIV-1 DNA. Nature 424, 94–98 (2003).

17. Browne, E.P., Allers, C. & Landau, N.R. Restriction of HIV-1 by APOBEC3G is cytidine deaminase-dependent. Virology 387, 313–321 (2009).

18. Miyagi, E. et al. Enzymatically active APOBEC3G is required for efficient inhibition of *Human immunodeficiency virus* type 1. J. Virol. 81, 13346–13353 (2007).

19. Schumacher, A.J., Hache, G., Macduff, D.A., Brown, W.L. & Harris, R. S. The DNA deaminase activity of human APOBEC3G is required for Ty1, MusD, and *Human immunodeficiency virus* type 1 restriction. J. Virol. 82, 2652–2660 (2008).

20. Sadler, H.A., Stenglein, M.D., Harris, R.S. & Mansky, L.M. APOBEC3G contributes to HIV-1 variation through sublethal mutagenesis. J. Virol. 84, 7396–7404 (2010).

21. Mulder, L.C., Harari, A. & Simon, V. Cytidine deamination induced HIV-1 drug resistance. Proc. Natl. Acad. Sci. USA 105, 5501–5506 (2008).

22. Russell, R.A., Moore, M.D., Hu, W.S. & Pathak, V.K. APOBEC3G induces a hypermutation gradient: purifying selection at multiple steps during HIV-1 replication results in levels of G-to-A mutations that are high in DNA, intermediate in cellular viral RNA, and low in virion RNA. Retrovirology 6, 16 (2009).

23. Hultquist, J.F. et al. Human and rhesus APOBEC3D, APOBEC3F, APOBEC3G, and APOBEC3H demonstrate a conserved capacity to restrict Vif-deficient HIV-1. J. Virol. 85, 11220–11234 (2011).

24. Zielonka J. et al. Restriction of *Equine infectious anemia virus* by equine APOBEC3 cytidine deaminases. J. Virol. 83, 7547–7559 (2011)

25. Delebecque F. et al. Restriction of *Foamy viruses* by APOBEC cytidine deaminases. J. Virol. 80, 605–614 (2006).

26. Mahieux, R. et al. Extensive editing of a small fraction of *Human T-cell leukemia virus* type 1 genomes by four APOBEC3 cytidine deaminases. J. Gen. Virol. 86, 2489–2494 (2005).

27. Dang, Y., Wang, X., Esselman, W.J. & Zheng, Y.H. Identification of APOBEC3DE as another antiretroviral factor from the human APOBEC family. J. Virol. 80, 10522–10533 (2006).

28. Bonvin, M. et al. Interferon-inducible expression of APOBEC3 editing enzymes in human hepatocytes and inhibition of *Hepatitis B virus* replication. Hepatology 43, 1364–1374 (2006).

29. Gee, P. et al. APOBEC1-mediated editing and attenuation of *Herpes simplex virus* 1 DNA indicate that neurons have an antiviral role during herpes simplex encephalitis. J. Virol. 2011 85, 9726–9736 (2011).

30. Suspène, R. et al. Genetic editing of *Herpes simplex virus* 1 and *Epstein-Barr herpesvirus* genomes by human APOBEC3 cytidine deaminases in culture and *in vivo*. J. Virol. 85, 7594–7602 (2011).

31. Fehrholz, M. et al. The innate antiviral factor APOBEC3G targets replication of measles, mumps and respiratory syncytial viruses. J. Gen. Virol. 93, 565–576 (2012).

32. Pauli, E.K. et al. High level expression of the anti-retroviral protein APOBEC3G is induced by influenza A virus but does not confer antiviral activity. Retrovirology 6, 38 (2009).

33. Wang, G. et al. APOBEC3F and APOBEC3G have no inhibition and hypermutation effect on the human *Influenza A virus*. Acta Virol. 52, 193–194 (2008).

34. Conticello, S.G., Thomas, C.J., Petersen-Mahrt, S.K. & Neuberger, M.S. Evolution of the AID/APOBEC family of polynucleotide (deoxy)cytidine deaminases. Mol. Biol. Evol. 22, 367–377 (2005).

35. Gowda, M. et al. *Magnaporthe grisea* infection triggers RNA variation and antisense transcript expression in rice. Plant Physiol. 144, 524–533 (2007).

36. Blow, M.J. et al. RNA editing of human microRNAs. Genome Biol. 7, R27 (2006).

37. Haas, M., Bureau, M., Geldreich, A., Yot, P. & Keller, M. *Cauliflower mosaic virus*: still in the news. Mol. Plant Pathol. 3, 419–429 (2002).

38. Suspène, R., Henry, M., Guillot, S., Wain-Hobson, S. & Vartanian, J.P. Recovery of APOBEC3-edited *Human immunodeficiency virus* G→A hypermutants by differential DNA denaturation PCR. J. Gen. Virol. 86, 125–129 (2005).

39. Chenault, K.D. & Melcher, U. Patterns of nucleotide sequence variation among *Cauliflower mosaic virus* isolates. Biochimie 76, 3–8 (1994).

40. Yu, Q. et al. Single-strand specificity of APOBEC3G accounts for minus-strand deamination of the HIV genome. Nat. Struct. Mol. Biol. 11, 435–42 (2004).

41. Marco, Y. & Howell, S. H. Intracellular forms of viral DNA consistent with a model of reverse transcriptional replication of the *Cauliflower mosaic virus* genome. Nucl. Acids Res. 12, 1517–1528 (1984).

42. Liddament, M.T., Brown, W.L., Schumacher, A.J. & Harris, R.S. APOBEC3F properties and hypermutation preferences indicate activity against HIV-1 *in vivo*. Curr. Biol. 14, 1385–91 (2004).

43. Kohli, R.M. et al. Local sequence targeting in the AID/APOBEC family differentially impacts retroviral restriction and antibody diversification. J. Biol. Chem. 52, 40956–40964 (2010).

44. Müller, V. & Bonhoeffer, S. Guanine-adenine bias: a general property of retroid viruses that is unrelated to host-induced hypermutation. Trends Genet. 21, 264–268 (2005).

45. Bendahmane, A., Querci, M., Kanyuka, K. & Baulcombe, D.C. *Agrobacterium* transient expression system as a tool for the isolation of disease resistance genes: application to the *Rx2* locus in potato. Plant J. 21, 73–81.

46. Schoelz, J.E., Shepherd, R.J. & Daubert, S. Region VI of *Cauliflower mosaic virus* encodes a host range determinant. Mol. Cell Biol. 6, 2632–2637 (1986).

47. Scholelz, J.E. & Shepherd, R.J. Host range control of *Cauliflower mosaic virus*. Virology 162, 30–37 (1988).

48. Schwab, R. et al. Highly specific gene silencing by artificial microRNAs in *Arabidopsis*. Plant Cell 18, 1121–1133 (2006).

49. Caddick, M.X. et al. An ethanol-inducible gene switch for plants used to manipulate carbon metabolism. Nat. Biotech. 16, 177–180 (1998).

50. Roslan, H.A. et al. Characterization of the ethanol-inducible *alc* gene-expression system in *Arabidopsis thaliana*. Plant J. 28, 225–235 (2001).

51. Clough, S.J. & Bent, A.F. Floral dip: a simplified method for *Agrobacterium*-mediated transformation of *Arabidopsis thaliana*. Plant J. 16, 735–743 (1998).

52. Martín, S. & Elena, S.F. Application of game theory to the interaction between plant viruses during mixed infections. J. Gen. Virol. 90, 2815–2820 (2009).

